# MODELING POLARITY SITE MOVEMENT IN YEAST VIA CALCIUM-MEDIATED REGULATION OF GAP ACTIVITY

**DOI:** 10.1101/2022.12.20.521256

**Authors:** Vatsal Parikh

## Abstract

Symmetry-breaking polarization is a process that occurs in many cells. In budding yeast cells specifically, this polarization is possible due to the presence of a positive feedback loop, which allows for one and only one front to develop. Furthermore, in mating yeast cells the detection of pheromones plays a crucial role in the cell’s development of a polarity site; however, due to the microscopic size of yeast cells, there is often error in the initial polarity site’s accuracy. As a result, yeast cells can correct their polarity site by allowing the GTP-Cdc42 cluster to “wander”. This is typically considered as the indecisive phase of a cell. Recent studies have identified that cytosolic Ca^2+^ bursts, caused by sexual pheromones, increase the activity of GTPase-activating protein (GAP). This project tests this from a modeling perspective by adding a module of code to an already established positive feedback model that allows periodic spikes of GAP activity to occur. The model was successfully able to simulate the effects of calcium-mediated GAP spikes by portraying oscillating polarity sites and “wandering” polarity sites, using specific parameters based on biochemical data.

## 1. Introduction

Cell polarity is a crucial aspect of how cells generate asymmetric distributions of molecules along the axis of a cell. Cells that are polarized are determined to have a “clear front-to-back axis” (Johnson et al., 2011). This spatial arrangement of molecules across the cell allows for various processes to occur such as “differentiation, localized membrane growth, activation of the immune response, directional cell migration, and vectorial transport of molecules across cell layers” (Drubin and Nelson, 1996). The budding yeast *Saccharomyces cerevisiae* is a popular model organism to study cell polarity. Budding yeasts are unicellular organisms which lack the ability to move in their environment (Wang et al., 2004). Recent studies have determined that the local accumulation of the Rho-family GTPases, polarity regulating proteins, determines the polarity site of a cell. The GTPase, Cdc42, recruits effector proteins which perform various downstream functions, including orientation of the actin cytoskeleton. This then allows for the cytoskeleton to be correctly positioned (Johnson et al., 2011).

It is understood that positive feedback plays an important role in polarity establishment, where GTP-Cdc42 promotes the activation of neighboring molecules of Cdc42 at the plasma membrane (Johnson et al., 2011). Current models utilize positive feedback to model yeast polarity development (Howell et al., 2012). Models are based on current genetic and biochemical data. These models incorporate Cdc42 and a Bem1-scaffolded complex which contains a PAK and the Cdc42-directed GEF (Howell et al., 2009). The positive feedback occurs since GTP-Cdc42 at the membrane recruits the cytoplasmic Bem1 complex, which in turn creates more active GTP-Cdc42 from the local inactive GDP-Cdc42 (Howell et al., 2012).

In yeast cells, G-protein-coupled-receptors (GPCRs) at the membrane guide directional growth and movement by interpreting signals outside of the cell and generating responses inside the cell (Insall et al., 2013). There are two types of haploid yeast cells of mating type: (MAT) **a** and MATα, which can mate with one another. For the yeast to be able to detect nearby mating partners the cells secrete pheromones that bind to the GCPRs of the opposing mating type cell. **a** cells can sense α-factor with sterile 2 [Ste2]. Likewise, α cells can sense **a**-factor with Ste3 (Wang et al., 2004). This sensing is crucial for *S. cerevisiae* to mate with other yeast cells. The accurate detection of extracellular gradients guides their development of polarity sites (Henderson et al., 2019).

However, due to the exceptionally small size of yeast cells (~4μm in diameter), high accuracy for spatial gradient sensing is challenging. Simulations have also confirmed this suspicion (Berg et al., 1977).

Recent studies have indicated that mating yeast cells can correct their initial error and move the polarity site based on pheromone gradients (Dyer et al., 2013; Hegemann et al., 2017; Kelley et al., 2015). At first, it seems unreasonable to assume that a Cdc42 cluster, which is constantly reinforced by positive feedback, would move. However, time-lapse imaging of yeast cells reveals that the Cdc42 cluster “wanders” around the cortex of the cell on a several-minute timescale (Dyer et al., 2013) (**Figure 1**).

**Figure 1.**
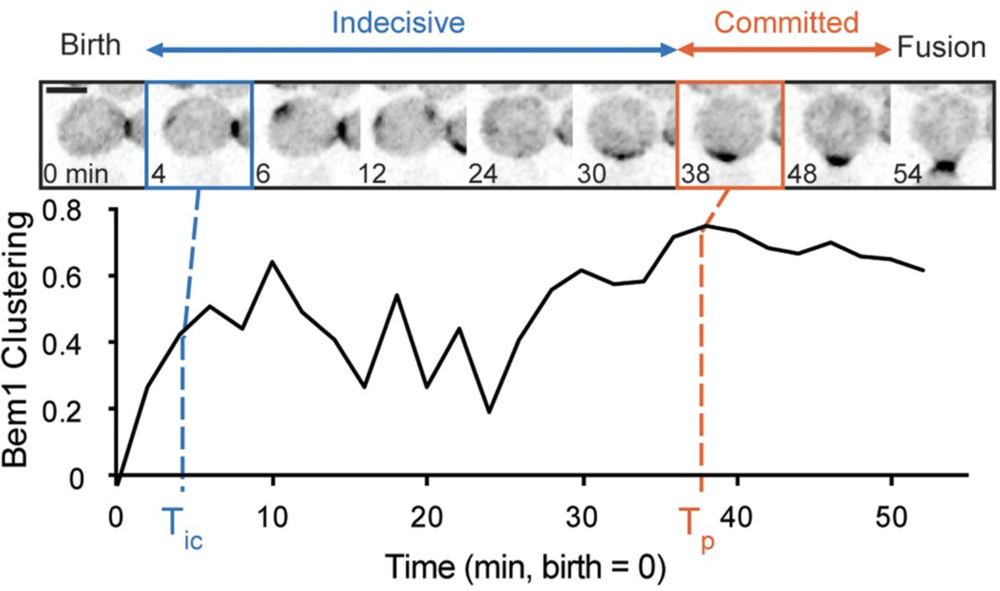
Development of polarity sites in mating yeast cells birth to fusion. Moments between birth and fusion include indecisive and committed periods of cell (Adapted from Henderson et al., 2019).

One hypothesis regarding the indecisive nature of polarity establishment is through transient and highly regulated spikes of cytosolic Ca^2+^+, which is already known to influence multiple cellular processes, and its impact on Rga2, a GTPase-activating protein (GAP), activity. Ca^2+^ signals are pervasive in cells since this cation can rapidly modify protein charge, shape, and function (Carbó et al., 2017). In mating yeast cells Ca^2+^ signals are a result of a response to environmental challenges such as the presence of mating pheromones (Ohsumi et al., 1985; Iida et al., 1990). Pheromone treated cells demonstrated noticeable responses to changes in pheromone concentrations with higher frequencies of Ca^2+^ bursts (Carbó et al., 2017). Furthermore, a connection between Ca^2+^ and GAP activity has been made since Rga2 has been identified as a calcineurin substrate, meaning calcineurin activates Rga2 and decreases Cdc42 signaling (Ly et al., 2017).

Although there are computational models that can model the development of polarity sites in yeast cells, there is yet a model that can model the recent discoveries made in yeast cells regarding their indecisive period. The aim of this paper is to see if there is a connection between the spikes in GAP activity, because of Ca^2+^ bursts, and the indecisive period in mating yeast cells observed by Henderson (Henderson et al., 2019). To do this the model will incorporate spikes in GAP activity to simulate the indecisive period observed in mating yeast cells.

## 2. Materials and Methods

### 2.1 Mathematical Modeling

The basis for the model used in this paper was adapted from a model for yeast cell polarity described in Howell et al. (2012). This model makes a few simplifying assumptions:

“(i) GDP-Cdc42 can exchange between the plasma membrane and cytoplasm. The membrane-bound and cytoplasmic forms are labeled as Cdc42D and Cdc42Ic, respectively. GTP-Cdc42 (Cdc42T) is always associated with the plasma membrane.
(ii) The Bem1 complex can exchange between cytoplasm (indicated as Bem1GEFc) and membrane (indicated as Bem1GEFm). Either form can bind to GTP-Cdc42 on the membrane, generating a complex indicated as Bem1GEFCdc42T.
(iii) The GEF activity of the Bem1 complex increases 2-fold when it binds GTP-Cdc42 (Howell et al., 2009).
(iv) GAP activity is spatially uniform and is incorporated in the first-order hydrolysis rate constant k2r.
(v) The GEF and GAP are not saturated by substrate (GDP-Cdc42 or GTP-Cdc42 respectively).
(vi) The cell dimensions, total Cdc42, and total Bem1 complex are all constant. The ratio of membrane volume to cytoplasmic volume is indicated by h.
(vii) All membrane-bound species have the same diffusion coefficient, Dm, and all cytosolic species the same diffusion coefficient, Dc, with Dc > > Dm” (Howell et al., 2012).

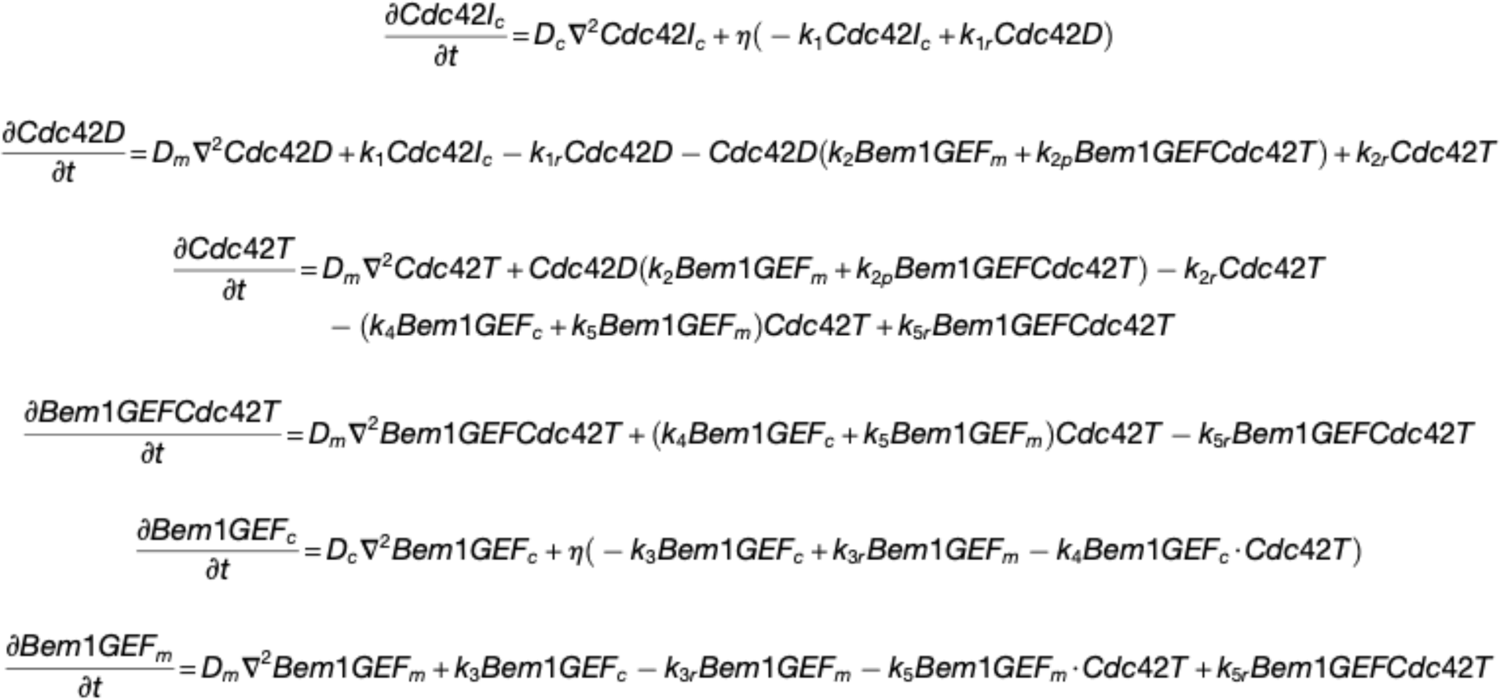

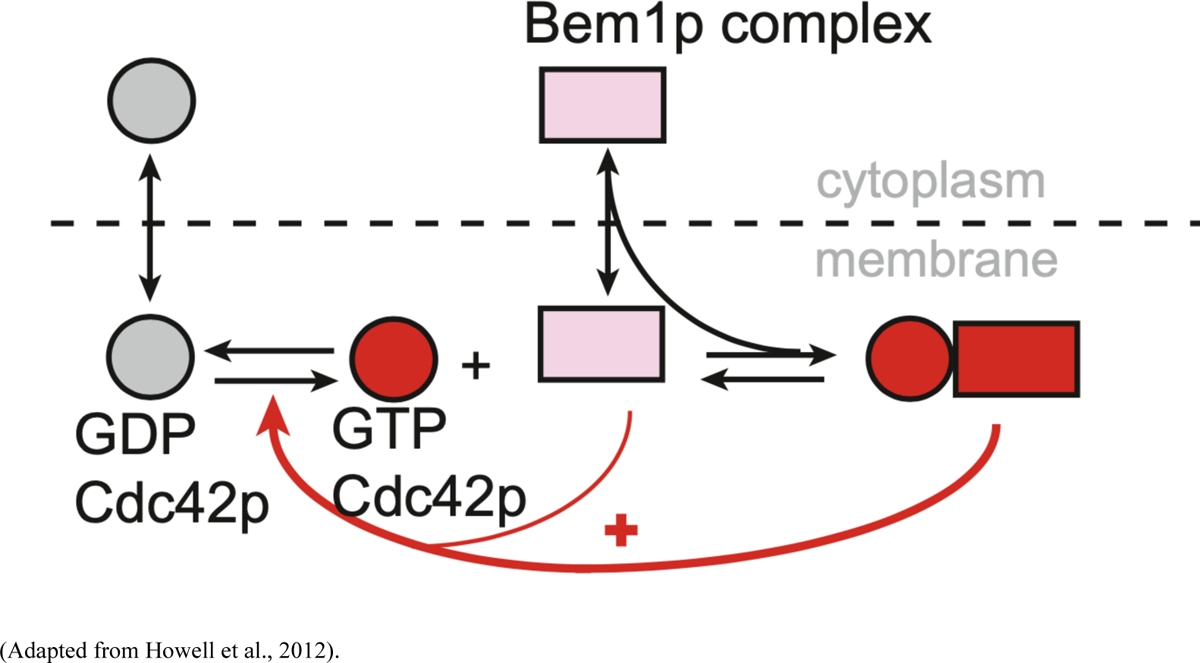

### 2.2 Modeling Calcium-Mediated GAP Spikes

It is suspected that bursts in cytosolic concentrations of Ca^2+^ during mating in yeast cells lead to transient increases in GAP activity through the action of calcineurin (Ly et al., 2017). It is also understood that these bursts occur at sporadic intervals for relatively short windows of time (Carbó et al., 2017) (**Figure 2**). To model the effects of transient GAP activation on polarity site behavior, I added a module to periodically alter: (1) the value of the k2r rate constant; (2) the duration of these “GAP spikes”; and (3) the frequency with which they occur.

**Figure 2.**
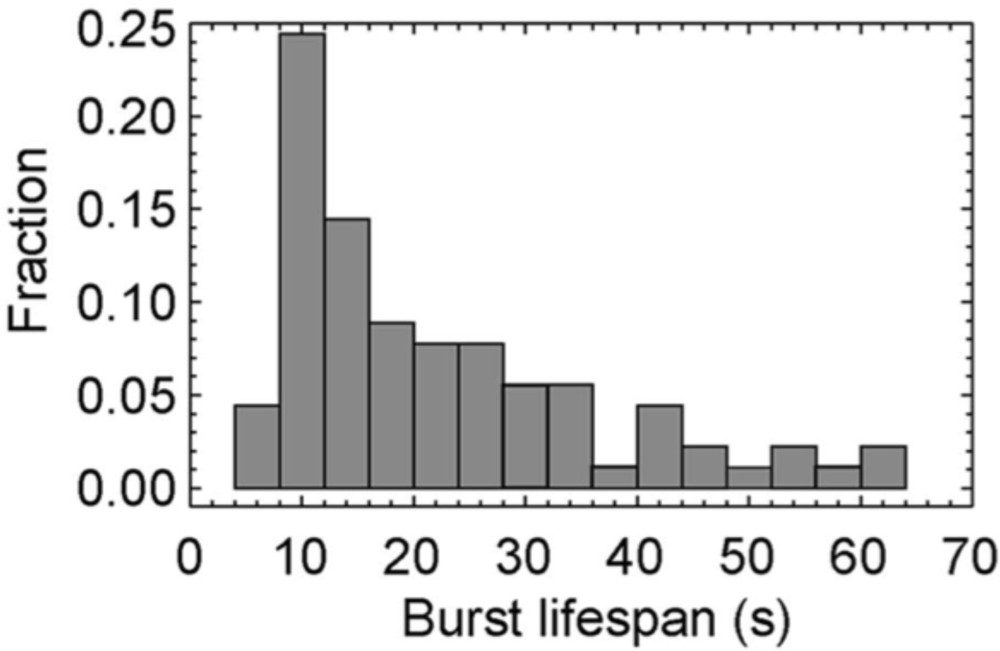
Various durations of Ca^2+^ bursts observed in mating yeast cells and frequency of their occurrence (Adapted from Carbó et al., 2017).

### 2.3 Identifying Reasonable Values for k2r Spikes

After incorporating logic to generate periodic spikes in the k2r rate constant, reasonable values needed to determine for the experiments. To determine this range, a single, pre-established polarity site was utilized (peak of GTP-Cdc42) at steady state as my initial condition. Various k2r values were then tested and the instantaneous impact on the polarity site was analyzed. Spike coefficients above 2.5 led to the disappearance of the polarity site within seconds, making those values infeasible and unrealistic. Through this testing it was determined that k2r values between 1.4 and 2.5 reduced the strength of the polarity patch enough to be noticeable, but not so much that the patch disappeared immediately at the beginning of the simulation.

### 2.4 Simulation and Analysis

The main parameters that were modified were the length of the k2r spikes, period between the spikes, and the strength of the spikes. Various combinations of these parameters were tested in simulations. Simulation, analysis, and visualization were performed in MATLAB R2022a.

## 3. Results and Discussion

### 3.1 Small GAP Spikes Induce Oscillating Polarity Sites

Various intervals and GAP spikes were simulated. These simulations had k2r values set such that they significantly lowered the GTP-Cdc42 peak, but not so much that it completely made the polarity site disappear (**Figure 3B**). The time intervals and periods were determined based on studies done on mating yeast cells identifying the time length of Ca^2+^ spikes. These periods of time were anywhere between 10 to 50 second bursts in GAP activity (Carbó et al., 2017). After the peak height was lowered, the k2r level was brought back to the level which allowed the cell to stay in steady state. Allowing this to occur, the GDP-Cdc42 begins to again convert back to GTP-Cdc42, and a peak begins to form again (**Figure 3C-D**). Furthermore, the peak reaches back to almost the same strength as the peak (**Figure 3E**). This is like what is seen in time-lapse imaging of cells with indecisive polarity sites. However, something that is different in the model to what is seen in the mating yeast cells is that if the patch clustering weakens it also begins to move along the cell cortex. This is not something that the model simulates as the polarity site pops up in the same location every time.

**Figure 3.**
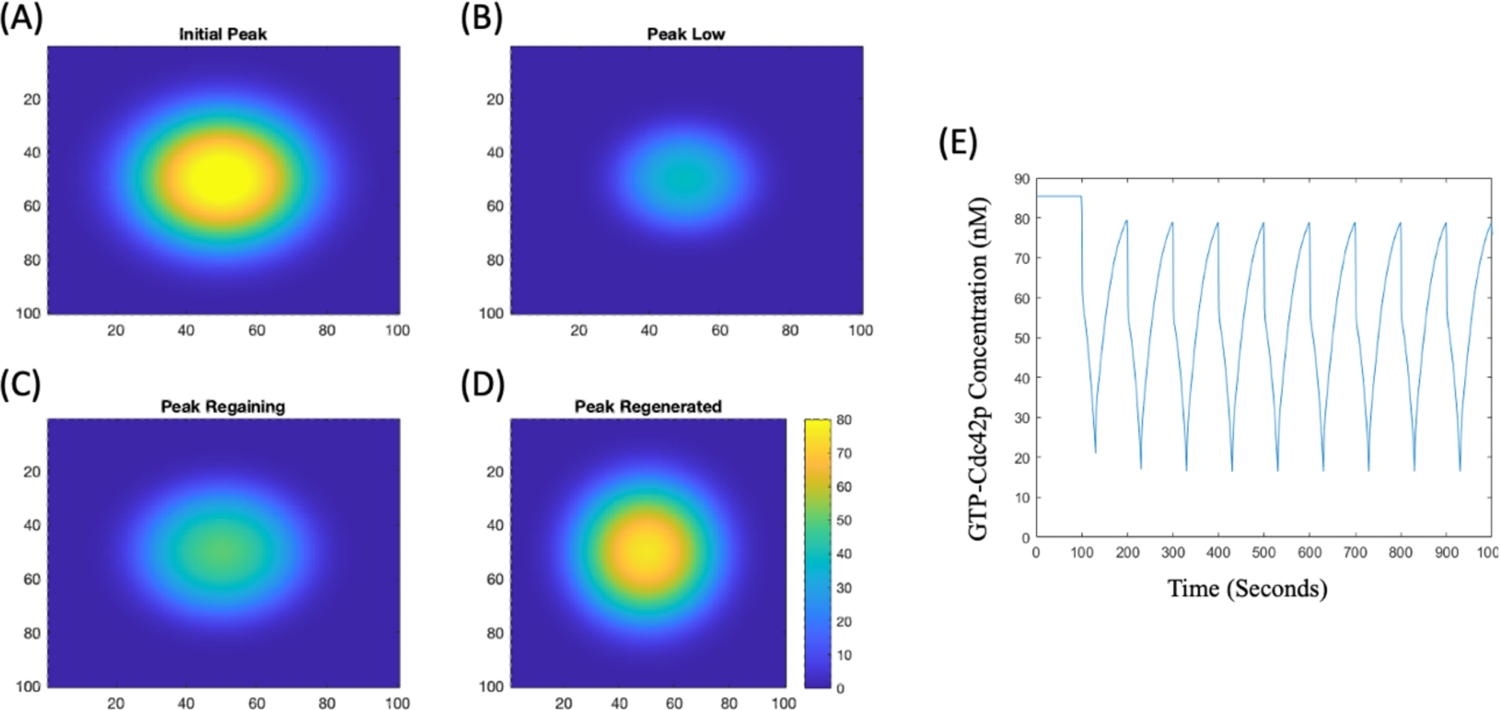
Plots of GTP-Cdc42 clustering in cell over time as calcium-mediated GAP spikes occur. **(A)** The strength of the initial peak before GAP spikes are introduced. **(B)** The strength of the peak at its lowest strength when the GAP spike is about to end. **(C)** Peak beginning to regenerate since GAP activity is back to steady state value. **(D)** The fully regenerated peak after having endured a GAP spike. **(E)** GTP-Cdc42 concentrations over time in cell as GAP spikes occur.

### 3.2 Regenerated Peak Begins to “Wander” When Original is Completely Diminished

Simulations were run in which the concentration of GTP-Cdc42 was completely depleted. This was done by setting the GAP activity to a level high enough to destroy the entire peak (**Figure 4B**). After the peak was completely destroyed, the k2r level was brought back to the level which allowed the cell to stay in steady state. Allowing this to occur, the GDP-Cdc42 begins to again convert back to GTP-Cdc42, and a peak begins to form again. However, two interesting things occur when this happens. The first is that the peak develops in a different location compared to the previous; this is also reliant on the fact that length of the GAP spike impacts how much the peak “wanders” (**Figure 4C**). This begins to mimic the “wandering” behavior seen in time-lapse imaging, where the peak completely disappears and reappears in another location on the cortex of the cell. Another observation seen from this “wandering” peak is that the concentration is nowhere near as strong as the original peak (**Figure 4D**). This is interesting to see since in time-lapse imaging it was seen that the concentration strength of GTP-Cdc42 was relatively similar for the various peaks formed in the cell. Further deterministic modeling studies could be done to see if more time or slight modifications in parameters after the peak is destroyed result in a strong cluster of GTP-Cdc42 forming in the cell. Additionally, another feature of the “wandering” peak is that it doesn’t completely disappear and reappear, but it rather starts to just move along the cortex of the cell and establishes in another part of the cell (Dyer et al., 2013; Hegemann et al., 2017; Kelley et al., 2015). A potential study for this aspect would be to reduce the strength of the peak enough so it is not completely established and see if it begins to wander as the Cdc42 begins to cluster again.

**Figure 4.**
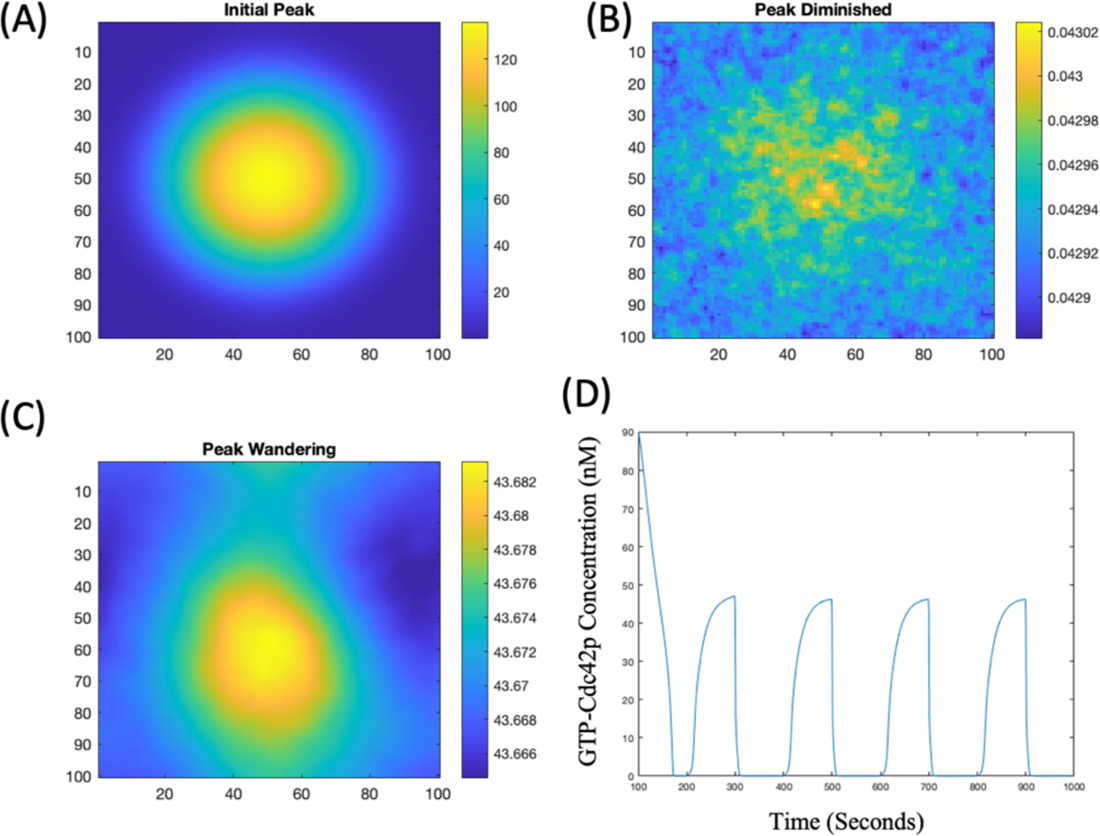
Plots of GTP-Cdc42 clustering over time as calcium-mediated GAP spikes completely destroy peaks. **(A)** A strong initial peak at steady state before GAP spikes are introduced. **(B)** No peak of GTP-Cdc42 after GAP spike. **(C)** New peak of GTP-Cdc42 is generated in a new location (now off-center) after the initial peak was diminished. **(D)** Rising and falling concentrations of GTP-Cdc42 over time as GAP spikes occur.

## 4. Acknowledgements

This project was done with the mentorship of Dr. Kyle Moran and the Lew Lab at Duke University. They not, however, authors of this paper.

## References

Berg HC, Purcell EM. Physics of chemoreception. Biophys J. 1977; 20(2):193–219. https://doi.org/10.1016/S0006-3495(77)85544-6 PMID: 911982; PubMed Central PMCID: PMC1473391.

Carbó N, Tarkowski N, Ipiña EP, Dawson SP, Aguilar PS. Sexual pheromone modulates the frequency of cytosolic Ca2+ bursts in Saccharomyces cerevisiae. Mol Biol Cell. 2017 Feb 15;28(4):501–510. doi: 10.1091/mbc.E16-07-0481. Epub 2016 Dec 28. PMID: 28031257; PMCID: PMC5305257.

Drubin DG, Nelson WJ. Origins of cell polarity. Cell. 1996 Feb 9;84(3):335–44. doi: 10.1016/s0092-8674(00)81278-7. PMID: 8608587.

Dyer JM, Savage NS, Jin M, Zyla TR, Elston TC, Lew DJ. Tracking shallow chemical gradients by actin-driven wandering of the polarization site. Curr Biol. 2013; 23(1):32–41. https://doi.org/10.1016/j.cub.2012.11.014 PMID: 23200992; PubMed Central PMCID: PMC3543483.

Hegemann B, Peter M. Local sampling paints a global picture: Local concentration measurements sense direction in complex chemical gradients. Bioessays. 2017; 39(7): 1600134. https://doi.org/10.1002/bies.201600134 PMID: 28556309.

Henderson NT, Pablo M, Ghose D, Clark-Cotton MR, Zyla TR, Nolen J, Elston TC, Lew DJ. Ratiometric GPCR signaling enables directional sensing in yeast. PLoS Biol. 2019 Oct 17;17(10):e3000484. doi: 10.1371/journal.pbio.3000484. PMID: 31622333; PMCID: PMC6818790.

Howell AS, Jin M, Wu CF, Zyla TR, Elston TC, Lew DJ. Negative feedback enhances robustness in the yeast polarity establishment circuit. Cell. 2012; 149(2):322–33. https://doi.org/10.1016/j.cell.2012.03. 012 PMID: 22500799; PubMed Central PMCID: PMC3680131.

Howell, A.S., Savage, N.S., Johnson, S.A., Bose, I., Wagner, A.W., Zyla, T.R., Nijhout, H.F., Reed, M.C., Goryachev, A.B., and Lew, D.J. (2009). Singularity in polarization: rewiring yeast cells to make two buds. Cell 139, 731–743.

Iida H, Yagawa Y, Anraku Y (1990). Essential role for induced Ca2+ influx followed by [Ca2+]i rise in maintaining viability of yeast cells late in the mating pheromone response pathway. A study of [Ca2+]i in single Saccharomyces cerevisiae cells with imaging of fura-2. J Biol Chem 265, 13391–13399.

Insall R. The interaction between pseudopods and extracellular signalling during chemotaxis and directed migration. Curr Opin Cell Biol. 2013; 25(5):526–31. https://doi.org/10.1016/j.ceb.2013.04.009 PMID: 23747069.

Johnson JM, Jin M, Lew DJ. Symmetry breaking and the establishment of cell polarity in budding yeast. Curr Opin Genet Dev. 2011; 21(6):740–6. https://doi.org/10.1016/j.gde.2011.09.007 PMID: 21955794; PubMed Central PMCID: PMC3224179.

Kelley JB, Dixit G, Sheetz JB, Venkatapurapu SP, Elston TC, Dohlman HG. RGS Proteins and Septins Cooperate to Promote Chemotropism by Regulating Polar Cap Mobility. Current Biology. 2015;25: 275–285

Ly N, Cyert MS. Calcineurin, the Ca2+-dependent phosphatase, regulates Rga2, a Cdc42 GTPase-activating protein, to modulate pheromone signaling. Mol Biol Cell. 2017 Mar 1;28(5):576–586. doi: 10.1091/mbc.E16-06-0432. Epub 2017 Jan 11. PMID: 28077617; PMCID: PMC5328617.

Ohsumi Y, Anraku Y (1985). Specific induction of Ca2+ transport activity in MATa cells of Saccharomyces cerevisiae by a mating pheromone, alpha factor. J Biol Chem 260, 10482–10486.

Wang Y, Dohlman HG. Pheromone signaling mechanisms in yeast: a prototypical sex machine. Sci-ence. 2004; 306(5701):1508–9. https://doi.org/10.1126/science.1104568 PMID: 15567849.

